# Efficient Reprogramming of the Epiblast Enables the Generation of Cloned Mice

**DOI:** 10.64898/2026.03.12.711217

**Authors:** Guomeng Li, Wandong Bao, Mei Hu, Heng Xie, Lianwei Li, Jianlin Xu, Shu Wei, Yue Teng, Yanyan Zhang, Yunan Chen, Lin Ran, Jiawen Liu, Juan Du, Muhammad Ameen Jamal, Zhanshan (Sam) Ma, Chikai Zhou, Jiangwei Lin

## Abstract

Somatic cell nuclear transfer (SCNT) can generate viable mammals despite pervasive epigenetic abnormalities in cloned embryos, yet the mechanism underlying this paradox remains unclear. Here we show, using single-cell transcriptomics, that efficient reprogramming occurs exclusively in the epiblast (EPI), but the not primitive endoderm (PrE) of late mouse SCNT blastocysts. Integrating this with previous findings of trophectoderm (TE) aberration, we propose the EPI as the sole effectively reprogrammed lineage. By innovatively employing a Lego-like multi-lineage embryonic aggregation approach, where the extra-embryonic lineages were replaced with fertilization-derived counterparts, we demonstrated that SCNT-derived EPI inherently possesses full-term developmental potential nearly identical to that of fertilized EPI (20.5% *vs*. 21.8% birth rate), functionally confirming its effective reprogramming. Our study uncovers a lineage-specific asymmetric reprogramming mode where the EPI specifically achieves effective reprogramming, thereby constituting the deterministic basis for cloned animal generation. This work also provides a versatile strategy for investigating lineage potency and function.

## Introduction

Somatic cell nuclear transfer (SCNT) represents a transformative biotechnology with profound implications for animal reproduction, endangered species preservation, and regenerative medicine^1–4^. Despite its successful application across various species, the efficiency of generating cloned animals remains notoriously low, largely attributed to the incomplete epigenetic reprogramming of the donor somatic nucleus^5,6^. Extensive research has established that SCNT embryos encounter formidable barriers immediately following nuclear transfer, including the resistance of somatic H3K9me3 to erasure, aberrant DNA methylation patterns, and failures in zygotic genome activation (ZGA)^7–10^. Consequently, it is widely accepted that cloned embryos harbor severe, genome-wide epigenetic aberrations that persist throughout pre-implantation development^11,12^. However, this presents a fundamental paradox: despite these pervasive epigenetic defects that theoretically compromise developmental competence, healthy cloned animals can still be successfully born^13–15^. This raises a fundamental question: is the birth of cloned animals merely a stochastic “game of chance” where rare cells accidentally escape epigenetic errors, or does an underlying deterministic mechanism exist that lays the foundation for cloned animal generation?

To decode this paradox, it is essential to examine the distinct cell lineages within the blastocyst, each of which possesses a specific developmental fate: the trophectoderm (TE), which forms the placenta; the primitive endoderm (PrE), which gives rise to the yolk sac; and the epiblast (EPI), which differentiates into the fetus proper^16,17^. Current research on reprogramming failure in SCNT blastocysts has predominantly focused on the extra-embryonic lineages, particularly the TE^18^. Indeed, our previous studies^19^, along with others, such as those highlighting the loss of H3K27me3-dependent non-canonical imprinting that leads to placental hyperplasia and early post-implantation lethality^12^, have demonstrated that the TE lineage in cloned embryos exhibits profound dysregulation. Given that the TE constitutes the majority of the blastocyst and is severely compromised, the key to understanding the “deterministic mechanism” of cloned animal generation likely lies within the inner cell mass (ICM). However, while studies have characterized global ICM defects^20^, current understanding of the precise molecular state, and thus the degree of reprogramming, of its two derived lineages, the PrE (extra-embryonic) and the EPI (embryonic), remains largely limited in SCNT blastocysts.

Here, we performed precise isolation of the ICM from late-stage SCNT blastocysts followed by single-cell RNA sequencing (scRNA-seq) to dissect the lineage-specific reprogramming status. Surprisingly, our analysis revealed a striking divergence in reprogramming fates: the PrE lineage displayed severe transcriptional aberrations, whereas the EPI lineage maintained a global gene expression profile that closely mirrored that of fertilization-derived (FD) controls. To functionally interrogate whether this molecularly “normal” EPI is indeed the driver of cloned animal birth, we developed an innovative “Lego-like” embryonic reconstruction strategy. By systematically replacing the potentially compromised SCNT extra-embryonic lineages (both PrE and TE) with their FD counterparts, we demonstrated that the EPI lineage from late-stage SCNT blastocysts possesses a developmental potential nearly identical to that of its natural counterpart, achieving a comparable full-term birth rate (20.5% *vs.* 21.8%). Collectively, our findings support the hypothesis that effective reprogramming occurs specifically in the EPI lineage of SCNT embryos. This specific reprogramming outcome, distinct from the global failures in extra-embryonic tissues, serves as the deterministic cellular foundation enabling the generation of cloned animals.

## Results

### ScRNA-seq reveals highly efficient reprogramming in the EPI lineage of late-stage mouse SCNT blastocysts

To investigate the reprogramming status of the ICM and its derived lineages, the PrE and EPI, in late-stage mouse SCNT blastocysts, we first profiled their gene expression landscapes. Using E4.5 blastocysts derived from the SCNT (NT) using cumulus cells and the fertilization-derived (FD) controls, we isolated ICMs by immunosurgery and performed bulk RNA sequencing (RNA-seq). Analysis of differentially expressed genes (DEGs) revealed that, overall, NT-ICMs exhibited distinct gene expression profiles compared to FD-ICMs, characterized by 515 up-regulated and 282 down-regulated DEGs (*p*.adjust < 0.05, Figure 1A). This finding is consistent with previous reports indicating global abnormalities in the ICM of cloned blastocysts^20^.

**Figure 1.**
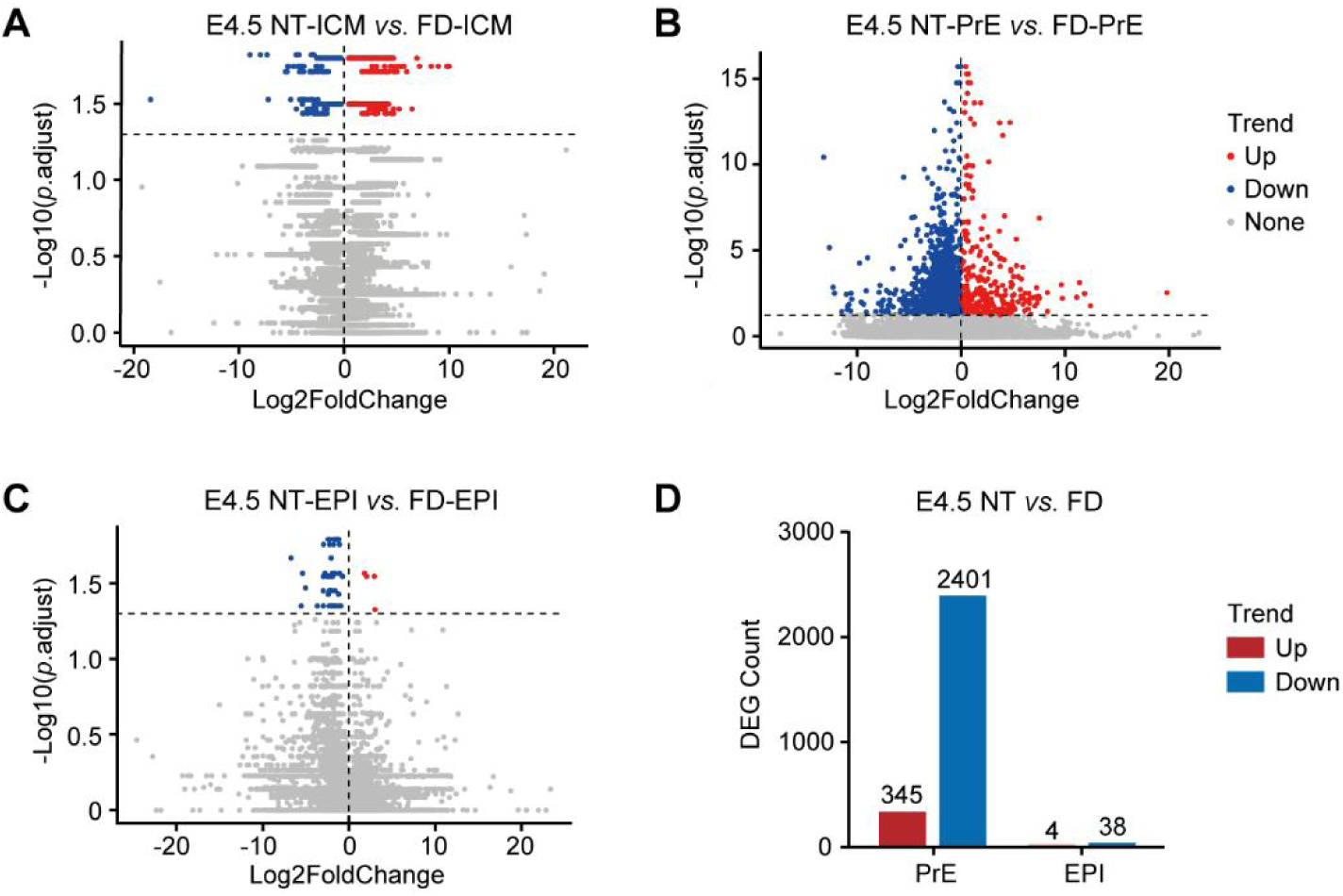
ScRNA-seq reveals highly efficient reprogramming in the EPI lineage of late-stage mouse SCNT blastocysts. (A) DEG analysis in E4.5 NT-ICM *vs*. FD-ICM based on their bulk RNA-seq (*p*.adjust < 0.05). (B, C) DEG analyses in NT-PrE *vs*. FD-PrE (B) and NT-EPI *vs*. FD-EPI (C) based on the scRNA-seq of E4.5 NT-ICMs and FD-ICMs (*p*.adjust < 0.05). (D) Statistics of DEG counts in E4.5 NT-PrE *vs*. FD-PrE and NT-EPI *vs*. FD-EPI.

For further dissecting the precise reprogramming states of the two constituent lineages, PrE and EPI, within late-stage cloned blastocysts, we dissociated E4.5 NT-ICMs (*n* = 3) and their FD counterparts (*n* = 2) into single cells (*n* = 84 and 53, respectively; Movie S1) and performed Smart-seq2 sequencing. Following quality control and clustering based on the expression of known lineage-specific marker genes, we obtained a total of 74 cells (comprising 63 PrE and 11 EPI cells) from NT blastocysts, and 49 cells (38 PrE and 11 EPI cells) from FD blastocysts (Figure S1A–D). We then performed DEG analyses in NT *vs.* FD (*p*.adjust < 0.05) separately for each lineage (Figure 1B, C). Strikingly, a massive number of DEGs were identified between NT-PrE and FD-PrE (*n* = 2,746, with 345 up-regulated and predominantly 2,401 down-regulated; Figure 1D), which showed significant associations with the biological processes (BPs) of kinase activity regulation, embryonic development, and RNA processing (Figure S2). In sharp contrast, only a minimal number of DEGs were detected between NT-EPI and FD-EPI (*n* = 42, with 4 up-regulated and also predominantly 38 down-regulated; Figure 1D). These results indicate that in late-stage cloned blastocysts, the PrE lineage suffers from severe reprogramming abnormalities, whereas the EPI lineage achieves highly efficient reprogramming at the transcriptome level.

Integrating these results with previous reports of severe TE dysregulation in cloned embryos^18,19^, we conclude that reprogramming failure in SCNT blastocysts is predominantly restricted to the extra-embryonic lineages (PrE and TE), while the EPI lineage, which develops the fetus proper, possesses the unique ability to achieve seemingly effective reprogramming. We thus hypothesize that this lineage-specific effective reprogramming endows the NT-EPI with the developmental competence to support full-term development, serving as the deterministic cellular foundation for the generation of cloned animals.

### Devising an induced lineage-based multi-lineage aggregation strategy to verify the hypothesis of SCNT-EPI developmental potential

To test this hypothesis, we designed a strategy to substitute the extra-embryonic lineages of NT blastocysts with their FD counterparts, retaining only the EPI lineage derived from SCNT. This approach aims to evaluate the developmental potential of the NT-EPI lineage for generating cloned mice by excluding the confounding effects of reprogramming abnormalities in the NT-TE and NT-PrE lineages. Initially, we planned to isolate NT-EPI and FD-PrE cells from E4.5 blastocysts – a stage when PrE and EPI lineages are considered fully differentiated and no longer interconvertible^21^ – and aggregate them with FD tetraploid (4N) embryos, which provide a functional TE lineage via tetraploid complementation^19,22^. The resulting reconstructed embryos (NT-EPI/FD-PrE/FD-4N), along with their strict controls (FD-EPI/FD-PrE/FD-4N), were intended to be transferred into surrogate females so that the developmental potential of the NT-EPI lineage could be measured by the birth rate of cloned mice. However, in practice, the PrE and EPI cells within the late-stage ICM are closely connected and thus difficult to precisely distinguish and separate, as confirmed in our experimental operations on the E4.5 blastocysts.

Nevertheless, it has been reported that treating early embryos with FGF4 and heparin (F4H) induces all E4.5 ICM cells to become Gata6-positive PrE cells^23^, while treatment with PD0325901 and CHIR99021 (two inhibitors of MEK and GSK3, respectively^24^; 2i) induces all E4.5 ICM cells to become Nanog-positive EPI cells^25^. Here, we refer to these cells as induced PrE (iPrE) and induced EPI (iEPI), respectively. Inspired by these findings, we sought to isolate iEPI or iPrE cells from E4.5 NT and FD blastocysts via induction followed by immunosurgery, and then aggregate them with FD-4N embryos. This alternative strategy allowed us to successfully obtain the desired “Lego-like” reconstructed embryos – specifically, NT-iEPI/FD-iPrE/FD-4N and their control FD-iEPI/FD-iPrE/FD-4N – thereby serving as a robust model to evaluate the capacity of the NT-EPI lineage to develop into cloned mice.

Before formally implementing this innovative reconstruction strategy, prudence dictated that we should comprehensively characterize these induced lineage cells to verify their identity and, importantly, confirm their equivalence to their naturally differentiated counterparts in both FD and NT contexts. To this end, we utilized FD and NT embryos carrying a compound transgenic background of *Nanog-GFP*^26^ and *Rosa26-CAG-tdTomato* (*R26-tdTomato*), which allows for the specific visualization of the pluripotent EPI lineage (GFP-positive) and the tracking of all donor-derived cells (tdTomato-positive), respectively. These embryos were treated with 2i or F4H from the 2-cell stage until the E4.5 blastocyst stage. The resulting induced ICMs (iICMs) – specifically, iEPI (2i-treated) or iPrE (F4H-treated) – were then isolated via immunosurgery and respectively aggregated with two 4-cell FD-4N embryos to monitor their chimeric behavior in the late-stage reconstructed blastocysts (Figure 2A).

**Figure 2.**
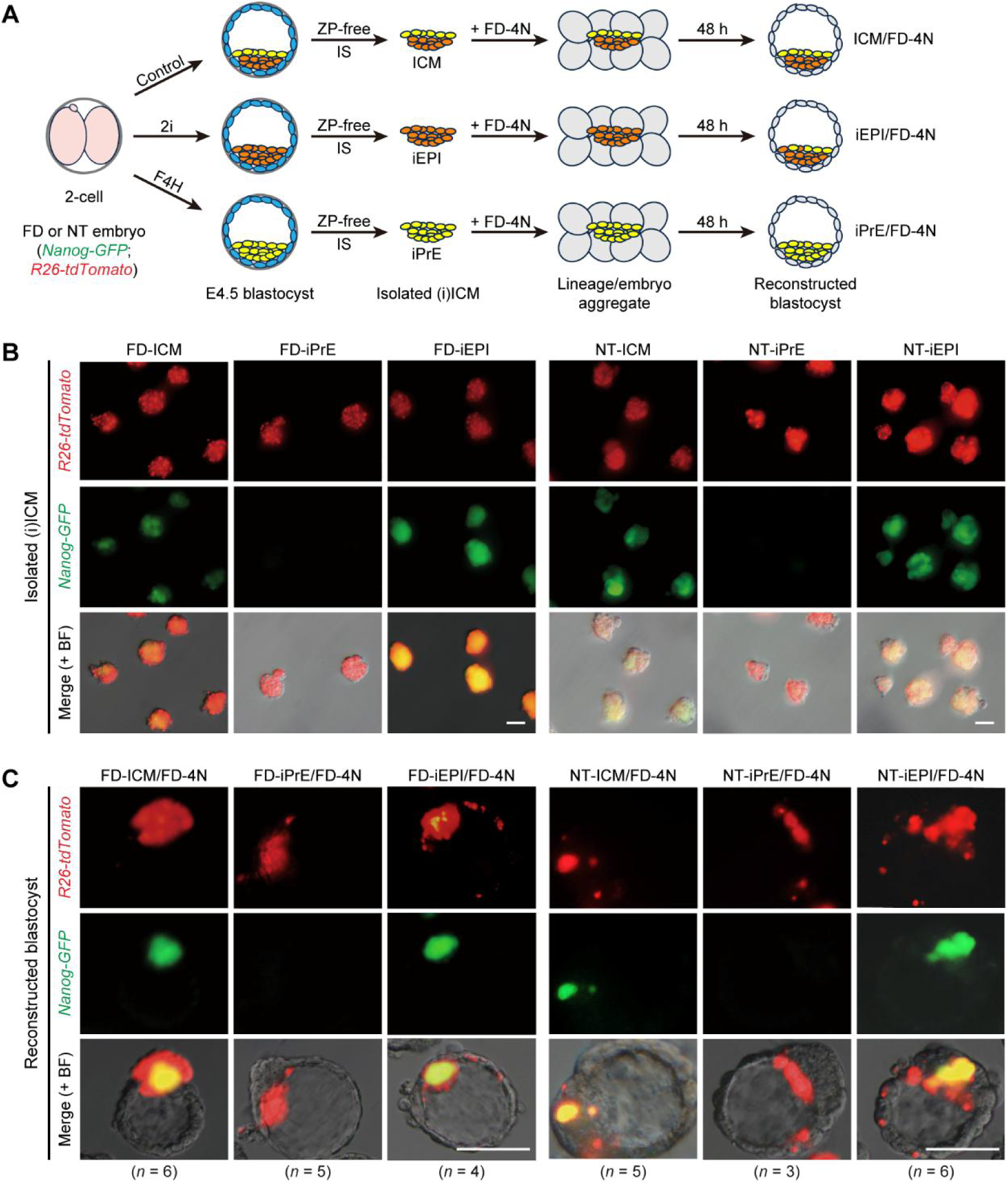
Chimeric behavior characteristics of induced lineages in reconstructed blastocysts under tetraploid complementation. (A) Schematic illustration of strategies for investigating chimeric behavior of induced lineages in reconstructed blastocysts under tetraploid complementation. The FD and NT embryos with *Nanog-GFP;R26-tdTomato* transgenic background were treated with 2i or F4H from the 2-cell stage until the E4.5 blastocyst stage; subsequently, the resulting iEPIs (2i-treated) or iPrEs (F4H-treated) were isolated via removal of zona pellucida and immunosurgery, and then aggregated with two 4-cell FD-4N embryos for detecting their chimeric behavior in the blastocysts reconstructed after 48-hour culture. The ICMs isolated from the naturally developed blastocysts were used as controls. ZP, zona pellucida; IS, immunosurgery. (B) Fluorescence microscopy of the (i)ICMs isolated from FD and NT blastocysts. Scale bar = 25 μm. (C) Fluorescence microscopy of the reconstructed blastocysts after 24-hour post-aggregation with FD-4N embryos for each (i)ICM type in (B). Scale bar = 65 μm.

First, we assessed the molecular identity of the induced lineages. Fluorescence microscopy of the isolated iICMs confirmed that *Nanog-GFP* expression was significantly repressed in both FD- and NT-iPrE, but maintained at high levels throughout the iEPIs (Figure 2B). Bulk RNA-seq analysis of these iICMs demonstrated that the iEPIs and iPrEs derived from both FD and NT embryos specifically enriched the marker gene expression of their corresponding lineages (Figure S3), validating the correctness of their induced lineage identities. Furthermore, based on our transcriptome data, no significant DEGs were identified between NT-iEPI and NT-EPI, or between FD-iEPI and FD-EPI (*p*.adjust < 0.05, Figure S4). This finding provides robust molecular evidence that the induced EPI lineages are equivalent to their naturally differentiated counterparts, thereby ensuring that using NT-iEPI *vs*. FD-iEPI as a surrogate for NT-EPI *vs*. FD-EPI as the sole experimental variable in our final aggregation strategy allows us to rigorously achieve our research objectives.

Next, to assess their integration capability, we aggregated each type of iICMs individually with two FD-4N embryos and cultured them *in vitro* for 48 hours to obtain late reconstructed blastocysts for monitoring the chimeric distribution. As expected, both FD-iPrE and NT-iPrE cells (tdTomato-positive only) were predominantly localized to the cavity side of the ICM in the reconstructed blastocysts – where the differentiated PrE lineage typically resides^16,27^ – and showed no *Nanog-GFP* expression (Figure 2C), indicating that the iPrEs maintained their lineage identity well within the reconstructed embryos. The chimeric behavior of both FD-and NT-iEPIs appeared slightly more complex: while the majority of cells (tdTomato-positive, GFP-positive) were identified as EPI cells embedded within the ICM region, a portion of cells (tdTomato-positive, GFP-negative) were found distributed in positions typical of PrE and TE (Figure 2C). This suggests that the iEPIs (isolated from E4.5 late blastocysts) exhibit some plasticity towards the two extra-embryonic lineages in this context. We speculate that this unusual lineage transdifferentiation may be attributed to the unique microenvironment formed within the reconstructed embryos generated by this specific lineage/embryo aggregation method, which might differ significantly from that of naturally developing embryos. Nonetheless, our data demonstrate that both iPrEs and iEPIs can efficiently integrate into the reconstructed embryos generated by lineage/embryo aggregation strategies and constitute their respective lineage components.

### The SCNT-derived EPI lineage achieves a low cloning efficiency via tetraploid complementation

To gain a preliminary understanding of the *in vivo* developmental potential of these induced lineages, particularly the NT-iEPI which is of our primary interest, we assessed their outcomes under tetraploid complementation following transfer into surrogate females. Given our observation that NT-iEPI cells can partially contribute to PrE and TE components in NT-iEPI/FD-4N reconstructed blastocysts, we initially hypothesized that these reconstructed embryos might possess a high potential for full-term development. To test this, NT embryos with a *CAG-GFP* transgenic background were processed to yield iEPIs (as described above), which were subsequently aggregated with FD-4N embryos. After 24 hours of *in vitro* culture, the aggregates formed blastocysts (Figure 3A, B) and were then transferred into surrogate females to evaluate their full-term developmental capacity.

**Figure 3.**
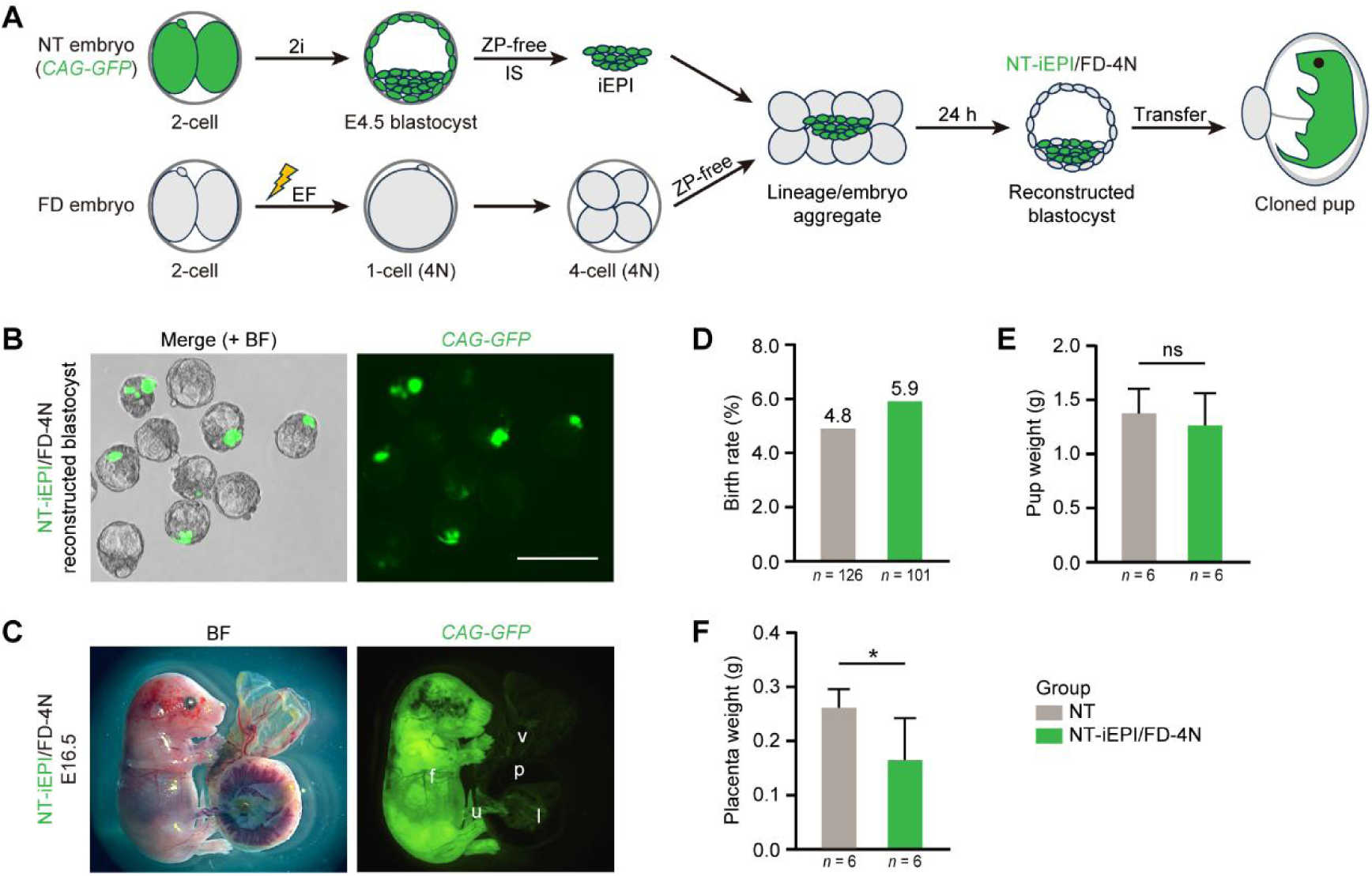
The SCNT-derived EPI lineage achieves a low cloning efficiency via tetraploid complementation. (A) Schematic illustration of strategy for evaluating the full-term development potential of NT-iEPI lineage based on tetraploid complementation. The NT embryo with *CAG-GFP* transgenic background was treated with 2i from the 2-cell stage until the E4.5 blastocyst stage; then, the resulting iEPI was isolated via removal of zona pellucida and immunosurgery, and aggregated with two 4-cell FD-4N embryos; after 24-hour *in vitro* culture, the reconstructed blastocyst developed from the aggregate was subsequently transferred to obtain cloned pup. ZP, zona pellucida; IS, immunosurgery; EF, electro-fusion. (B) Fluorescence microscopy of the NT-iEPI/FD-4N reconstructed blastocysts after 24-hour *in vitro* culture post-aggregation. Scale bar = 100 μm. (C) Fluorescence microscopy of the E16.5 fetus and extra-embryonic tissues generated by NT-iEPI/FD-4N strategy. f, fetus; u, umbilical cord; v, visceral yolk sac; l, labyrinth zone; p, placenta. (D–F) Statistics of the birth rate (D), body weight (E), and placenta weight (F) of the cloned pups generated by NT-iEPI/FD-4N strategy. The NT group was used as a baseline control. *, *p* < 0.05; ns, not significant; Student’s t-test.

Transgenic fluorescence imaging at E16.5 revealed that the fetuses generated from NT-iEPI/FD-4N were ubiquitously GFP-positive (Figure 3C), confirming their origin entirely from the NT-iEPI lineage. Meanwhile, GFP signal was also observed in the extra-embryonic tissues, such as the umbilical cord, visceral yolk sac, and the labyrinth zone of the placenta (Figure 3C), consistent with the reported involvement of EPI lineage in the development of these structures^28,29^. However, no detectable GFP signal was found in any other area of the placenta (Figure 3C), standing in contrast to the phenomenon observed in NT-iEPI/FD-4N blastocysts, where the NT-iEPI cells showed a tendency to transdifferentiate into the extra-embryonic lineages including the TE (Figure 2C) that is responsible for forming the placenta^30^. Ultimately, we transferred a total of 101 NT-iEPI/FD-4N reconstructed blastocysts and merely obtained 6 live cloned pups, representing a birth rate of 5.9%. This efficiency was only marginally higher than that of the baseline NT control group (4.8%, 6/126; Figure 3D, Table 1), in which NT embryos were cultured to the blastocyst stage *in vitro* and then transferred. The body weight of the cloned pups generated from NT-iEPI/FD-4N showed no significant difference compared to the NT group (Figure 3E), whereas their placental weight was significantly lower (Figure 3F), which is likely attributed to the rescue of SCNT-associated placental hyperplasia by the FD-TE lineage provided via tetraploid complementation^19,20^.

**Table 1.**
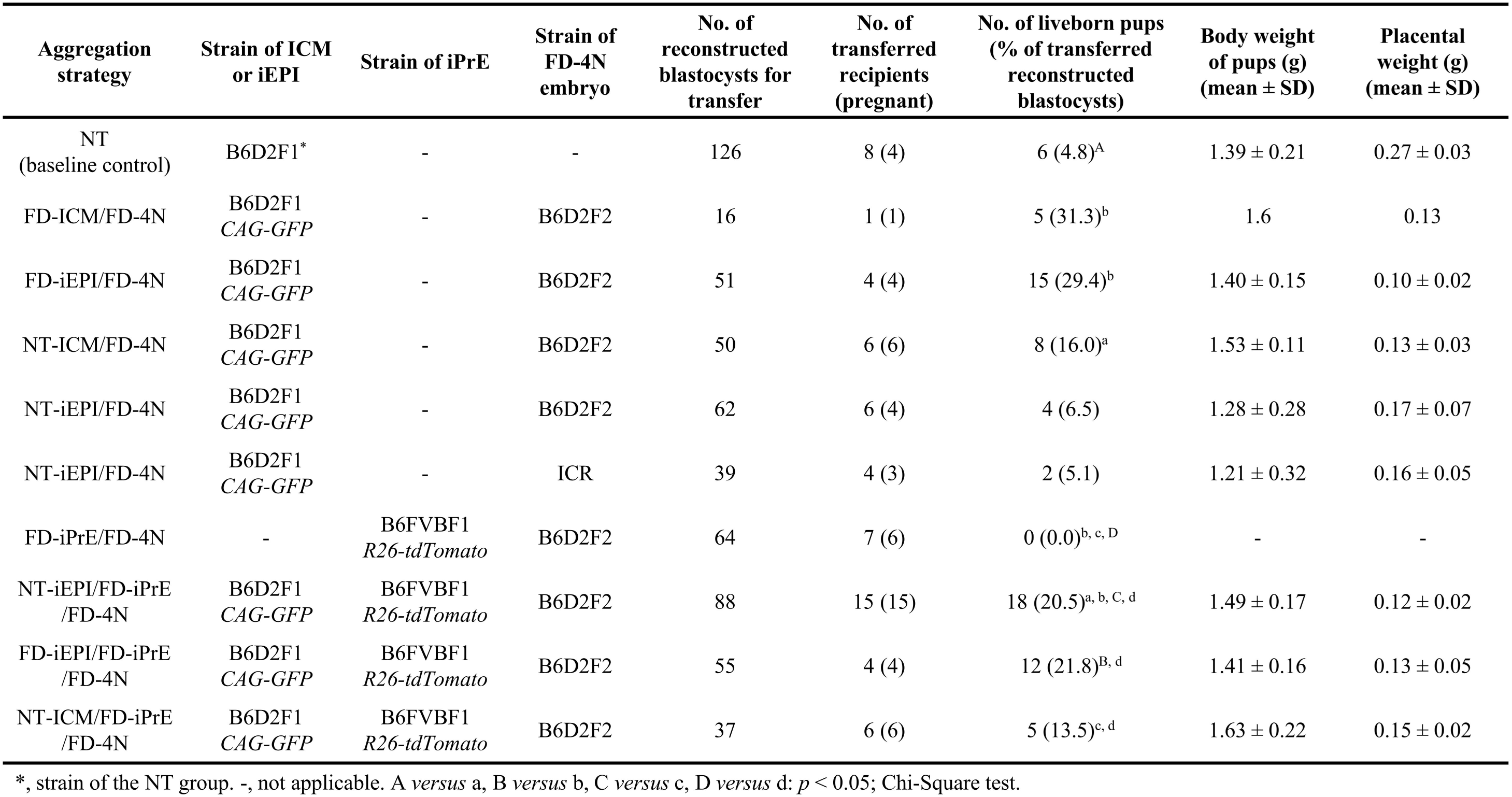
Statistical analysis of *in vivo* developmental outcomes of reconstructed embryos generated by various lineage/embryo aggregation strategies.

Here, we speculate that the *de novo* NT-TE cells, as well as NT-PrE cells, derived from the NT-iEPI might still harbor severe reprogramming abnormalities due to the lack of maternal epigenetic inheritance (e.g., H3K27me3^10,11,31^), and consequently, act as functionally defective cells that are gradually outcompeted and replaced by the non-fluorescent, FD-4N-derived extra-embryonic lineages during subsequent development. If this is indeed the case, such futile lineage transdifferentiation by the NT-iEPI likely represents a detrimental drain on the cell pool destined for the fetus proper, which might even be exacerbated by a persistent lack of effective negative feedback due to the failure of transdifferentiation into functional extra-embryonic lineages, especially its critical neighbour, the PrE lineage. In support of this, immunofluorescence staining revealed that within the ICM regions of numerous NT-iEPI/FD-4N reconstructed blastocysts, the NT-iEPI(-derived) cells, despite exhibiting *Nanog* expression, almost universally displayed significant *Gata6* expression (Figure S5), suggesting an aberrantly high degree of ongoing transdifferentiation toward the PrE lineage. This would therefore drastically diminish the NT-iEPI cell pool available for embryonic proper development, which helps to explain the severely compromised success rate of generating full-term cloned pups under this aggregation strategy.

In parallel, utilizing the same methodology, we investigated the birth efficiency of FD-iEPI, NT-ICM, and FD-ICM aggregated with FD-4N embryos (Table 1). Possibly owing to its purely FD cellular composition and the consequent potential to generate a relatively normal PrE lineage, the FD-iEPI/FD-4N group achieved a remarkably high birth rate of 29.4% (15/51), which was only marginally lower than the 31.3% (5/16) observed in the FD-ICM/FD-4N group. In contrast, the NT-ICM/FD-4N group yielded a relatively lower birth rate of 16.0% (8/50), theoretically constrained by the overall SCNT origin of its ICM. However, this rate was still notably higher than the 5.9% observed in the NT-iEPI/FD-4N group, suggesting that although the NT-PrE lineage harbors severe reprogramming abnormalities, its presence (as in NT-ICM/FD-4N) is still more advantageous than its initial absence (as in NT-iEPI/FD-4N) for facilitating the successful birth of cloned pups to some extent.

Additionally, we utilized *R26-tdTomato* FD embryos to derive iPrE cells and tested their developmental potential *in vivo* via tetraploid complementation (Figure S6A). As expected, following the transfer of blastocysts developed from FD-iPrE/FD-4N aggregates (Figure S6B), no live pups were obtained (0%, 0/64; Table 1), with only occasional resorption sites observed upon dissection (data not shown). This failure should be attributed to the lack of an embryonic (EPI) lineage competent to form the fetus proper in FD-iPrE/FD-4N embryos, where the FD-4N component can only effectively support extra-embryonic development^19,20^ and the FD-iPrE was observed to strictly maintain its lineage identity (Figure 2C) as also further confirmed by immunofluorescence assays (Figure S6C). Given these results, the NT-iPrE appeared even less likely to support full-term development via tetraploid complementation, further due to its potential SCNT-associated reprogramming defects; thus, *in vivo* developmental testing was omitted for this group.

### The SCNT-derived EPI lineage possesses full-term developmental potential equivalent to that of the fertilization-derived EPI lineage

Having comprehensively characterized the induced lineages in terms of their molecular signatures, chimeric integration capabilities, and *in vivo* developmental potentials under tetraploid complementation, we proceeded to the definitive phase of our study by implementing the “Lego-like” multi-lineage embryonic aggregation strategy, aiming to rigorously explore the developmental potential of the SCNT-derived EPI lineage for generating full-term cloned mice under conditions where the extra-embryonic lineages are fully functional and normal. Using *CAG-GFP* NT embryos and *R26-tdTomato* FD embryos, we induced and isolated iEPI and iPrE cells, respectively, as described above, and subsequently aggregated them simultaneously with two FD-4N embryos. After 24 hours of *in vitro* culture, the aggregates developed into NT-iEPI/FD-iPrE/FD-4N reconstructed blastocysts, which were then transferred into surrogate females to generate cloned pups (Figure 4A).

**Figure 4.**
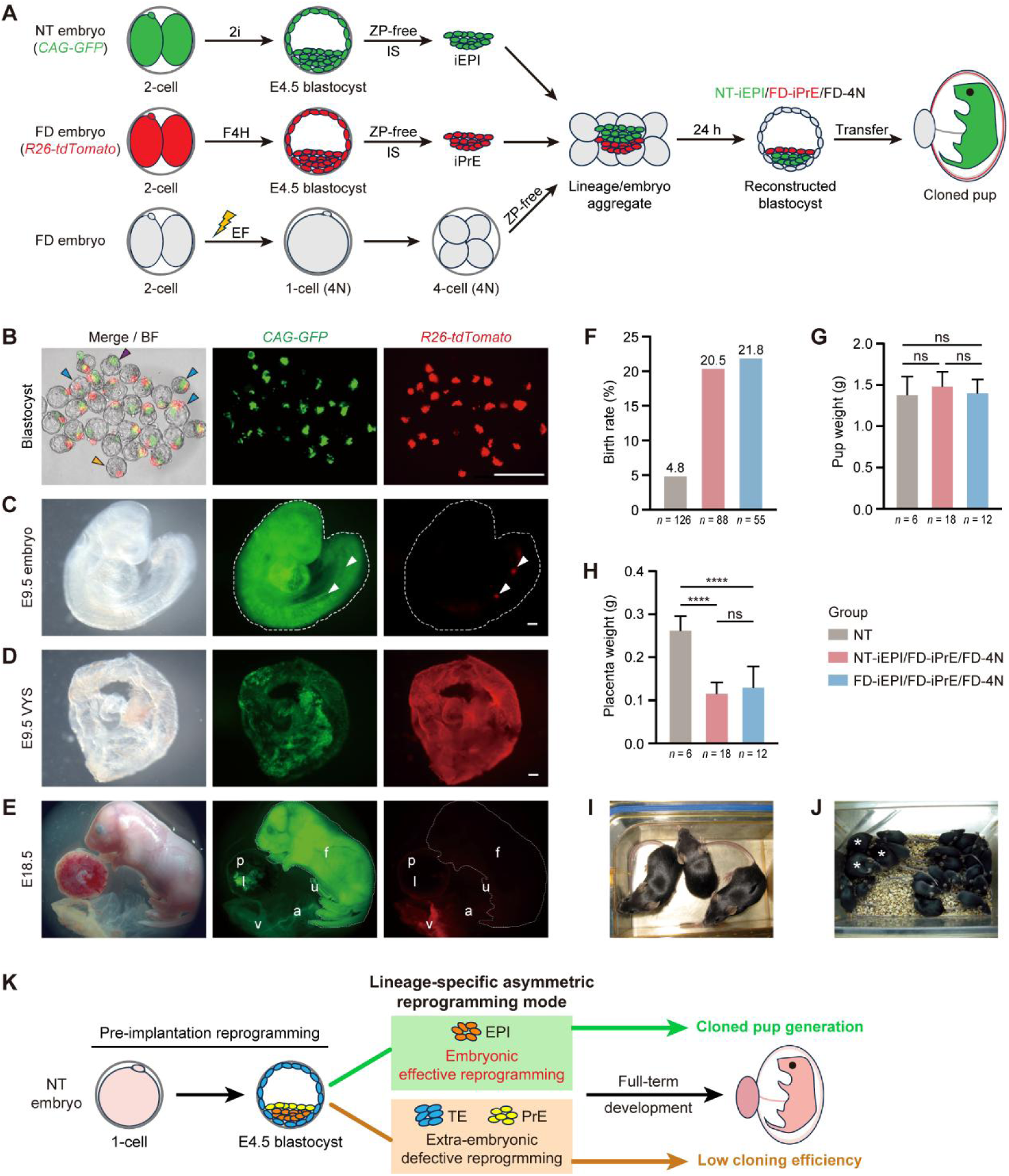
Tri-lineage aggregation strategy validates the inherent robust full-term developmental potential of SCNT-derived EPI lineage. (A) Schematic illustration of strategy for evaluating the full-term development potential of NT-iEPI lineage based on tri-lineage embryonic aggregation. Specifically, the *CAG-GFP* NT embryo and *R26-tdTomato* FD embryo were treated respectively with 2i and F4H from the 2-cell stage until the E4.5 blastocyst stage; then, the resulting NT-iEPI and FD-iPrE were isolated via removal of zona pellucida and immunosurgery, and aggregated with two 4-cell FD-4N embryos; after 24-hour *in vitro* culture, the reconstructed blastocyst developed from the aggregate was subsequently transferred to obtain cloned pup. ZP, zona pellucida; IS, immunosurgery; EF, electro-fusion. (B–E) Fluorescence microscopy of the reconstructed blastocysts after 24-hour *in vitro* culture post-aggregation (B), E9.5 embryo (C) and visceral yolk sac (D), and E18.5 fetus and extra-embryonic tissues (E) generated by NT-iEPI/FD-iPrE/FD-4N strategy. The arrowheads in (B) indicate the representative reconstructed blastocysts with significant dual fluorescence (blue arrowheads) that were selected for subsequent transfer operation, and the ones with residual single fluorescence (tdTomato only, yellow arrowhead; GFP only, purple arrowhead) that were thereafter discarded. The white dashed lines indicate the outline of the E9.5 embryo (C) and E18.5 fetus (E). The white arrowheads indicate the significant tdTomato-positive positions in E9.5 embryo (C). f, fetus; u, umbilical cord; v, visceral yolk sac; a, amnion; l, labyrinth zone; p, placenta (E). Scale bar = 100 μm (B), 2.5 mm (C, D). (F–H) Statistics of the birth rate (F), body weight (G), and placenta weight (H) of the neonatal pups generated by the tri-lineage aggregation strategies. The NT group was used as a baseline control. ****, *p* < 0.0001; ns, not significant; Student’s t-test. (I) Adult Tri-NT mice. (J) Tri-NT mice were fertile to produce healthy offspring. The Tri-NT female parents are indicated (*). (K) Theoretical model of “lineage-specific asymmetric reprogramming” in SCNT-derived late blastocyst. As pre-implantation reprogramming of NT embryos progresses to the late blastocyst stage, the differentiated cell lineages exhibit an asymmetric reprogramming mode: the embryonic lineage (EPI) achieves effective reprogramming to exercise fully functional pluripotency, constituting the biological foundation of cloned pup generation, whereas the extra-embryonic lineages (TE and PrE) display seriously defective reprogramming, leading to extra-embryonic dysfunction and consequent low cloning efficiency.

Transgenic fluorescence imaging revealed that approximately 75% of the reconstructed blastocysts developed from the aggregates retained significant amounts of both NT-iEPI and FD-iPrE components (GFP-positive, tdTomato-positive; Figure 4B), and these were selected for subsequent transfer. Immunofluorescence staining showed that within the ICM regions of reconstructed blastocysts, the FD-iPrE(-derived) cells (tdTomato-positive) exhibited *Gata6* expression, whereas the NT-iEPI(-derived) cells (GFP-positive) were uniformly negative (Figure S7), indicating no transdifferentiation from the NT-iEPI to PrE lineage in this context. Subsequently, we examined the fluorescence distribution at E9.5 and found that the fetus proper was almost exclusively GFP-positive (derived from NT-iEPI), with only a few tdTomato-positive cells (derived from FD-iPrE) observed in the gut region, particularly the hindgut (Figure 4C). This distribution aligns perfectly with current knowledge regarding EPI and PrE lineage contributions during embryonic development, where the EPI forms the fetus proper^32–34^, while a minority of PrE cells transiently contribute to parts of the gut endoderm^35,36^. The VYS enveloping the E9.5 fetus exhibited double positivity for GFP and tdTomato (Figure 4D), consistent with the fact that the VYS is composed of an inner layer of extra-embryonic mesoderm derived from EPI and an outer layer of visceral endoderm derived from PrE^37,38^. Furthermore, this dual-fluorescence pattern was also observed in the VYS obtained from cesarean sections at E18.5 (Figure 4E). At this stage, prominent GFP signals were accurately detected in other structures that EPI developmentally involves, including the fetus proper, amnion, umbilical cord, and the labyrinth zone of the placenta^28,29,39^ (Figure 4E), with the majority of the placental tissue being non-fluorescent, presumably originating from the FD-4N embryos. These data robustly demonstrate that in our multi-lineage aggregation strategy, both NT-iEPI and FD-iPrE strictly adhere to their respective lineage identities and correctly fulfill their developmental fates.

Ultimately, we transferred a total of 88 NT-iEPI/FD-iPrE/FD-4N reconstructed blastocysts and successfully obtained 18 live cloned pups (designated as Tri-NT mice due to their generation via tri-lineage/embryo aggregation), achieving a birth rate of 20.5% (Figure 4F, Table 1). This rate was significantly higher than the 4.8% (6/126) of the baseline NT control group, indicating that the holistic replacement of extra-embryonic lineages with their normal counterparts can drastically improve cloning efficiency. Crucially, compared to the strict control group FD-iEPI/FD-iPrE/FD-4N prepared using the same methodology, the birth rate of Tri-NT mice was nearly identical to that of the control pups (designated as Tri-FD mice), which stood at 21.8% (12/55; Figure 4F, Table 1). This provides direct evidence that NT-iEPI (equivalent to NT-EPI) possesses a full-term developmental potential comparable to that of FD-iEPI (equivalent to FD-EPI). Moreover, Tri-NT pups showed no significant differences in body weight (Figure 4G) or placental weight (Figure 4H) compared to Tri-FD pups. Attributable to tetraploid complementation, the placental weight of Tri-NT pups was significantly ameliorated compared to the NT group (Figure 4H), so was the morphology of the labyrinth and spongiotrophoblast layers within their placentae, as indicated by histologic sectioning (Figure S8). The Tri-NT mice could grow to adulthood (Figure 4I) and were fertile to produce healthy offspring (Figure 4J). Furthermore, to confirm the reproducibility of these findings, we performed an independent replicate of the experiments, which yielded statistical outcomes that recapitulated the trends presented above (Figure S9A–C).

Compared to the NT-iEPI/FD-4N aggregation, the NT-iEPI/FD-iPrE/FD-4N strategy achieved a substantial increase in cloning efficiency (20.5% *vs*. 5.9%), attributed to the inclusion of FD-iPrE. The underlying mechanism likely involves two factors: first, the presence of FD-iPrE at the onset of aggregation allows the NT-iEPI to maintain a more abundant cell pool for fetal development by eliminating its need for compensatory transdifferentiation into the PrE lineage; second, as a representative of the normal PrE lineage, FD-iPrE theoretically supports NT-iEPI embryonic development more effectively than the potentially dysfunctional *de novo* NT-PrE cells generated in the NT-iEPI/FD-4N strategy. Therefore, only by incorporating FD-iPrE into the aggregation can the developmental potential of NT-iEPI to form the embryo proper and achieve full-term development be realized to the maximum extent, confirming the feasibility and necessity of implementing the NT-iEPI/FD-iPrE/FD-4N strategy to achieve the objectives of this study.

In summary, by implementing this innovative multi-lineage embryonic aggregation strategy to holistically replace the reprogramming-compromised extra-embryonic lineages with their FD counterparts, we successfully unlocked the latent full-term developmental potential of the NT-(i)EPI lineage. Our data demonstrate that the NT-(i)EPI lineage inherently possesses developmental competence equivalent to that of the FD-(i)EPI lineage, thereby serving as the core driver for the generation of cloned mice. Finally, based on our findings and existing studies, we propose a theoretical model of “lineage-specific asymmetric reprogramming” governing the developmental potential of cloned embryos: while SCNT-induced reprogramming defects are pervasively retained in the extra-embryonic lineages (TE and PrE), severely compromising the cloning efficiency, the embryonic lineage (EPI) specifically achieves effective reprogramming and possesses intact pluripotency, thereby constituting the biological foundation that enables the generation of cloned animals (Figure 4K).

## Discussion

Historically, the prevailing consensus regarding reprogramming in SCNT embryos posits that the failure to fully surmount early epigenetic barriers, such as the resistance of H3K9me3 erasure^7,40,41^ that can lead to defective ZGA^5,42^, leaves cloned embryos burdened with widespread and severe reprogramming defects across the entire genome^4,7^. However, this perception of “global failure” has failed to coherently reconcile a core paradox: how can SCNT embryos, theoretically laden with such a profound and pervasive epigenetic burden, still possess the probability to break through these barriers and develop into completely healthy cloned animals? This cognitive gap largely stems from the fact that prior research has predominantly focused on the global embryonic state following nuclear transfer^11,43^, while overlooking the potential divergence in reprogramming fates among distinct cell lineages as development progresses to the blastocyst stage. In this study, by precisely dissecting the fine architecture within ICM of late-stage blastocysts and coupling scRNA-seq analysis with functional reconstruction strategies, we have uncovered and confirmed a mode of “lineage-specific asymmetric reprogramming”. It is crucial to emphasize that, for those cloned embryos capable of successfully developing to the late blastocyst stage, reprogramming failure is not persistently diffuse throughout the embryo. Instead, our findings reveal that defects are primarily restricted within the extra-embryonic lineages (TE and PrE); in sharp contrast, the EPI lineage, destined to form the fetus, specifically exhibits a remarkably effective reprogrammed state at this stage. This discovery not only theoretically helps to resolve the aforementioned “survivor paradox” but also offers a fundamental revelation: the birth of cloned animals is not the result of absolutely stochastic chance, but rather is founded upon a deterministic biological mechanism – the successful reconstruction of intact pluripotency specifically within the EPI lineage during the critical window of blastocyst formation.

The lineage-specific effective reprogramming observed in SCNT blastocysts might be intrinsically linked to the developmental properties of the EPI lineage. Accumulating evidence indicates that maternal H3K27me3-dependent non-canonical imprinting predominantly resides and functions within the extra-embryonic lineages, as exemplified by its critical role in ensuring the activity of the maternal X chromosome by repressing *Xist*^44^ as well as in regulating the expression of key developmental genes^12^, whereas these imprints are generally transient or dispensable in the EPI lineage^45^. Owing to the inherent lack of these oocyte-specific H3K27me3 modifications in SCNT donor nuclei, the extra-embryonic lineages of cloned embryos are thus foreseeable to suffer from severe and potentially irreparable gene expression dysregulation. By contrast, the EPI lineage, due to its low dependency on these imprints, appears relatively insensitive to this specific epigenetic defect, thereby exhibiting a markedly superior reprogramming state. Nevertheless, the EPI lineage of cloned embryos might be more than merely a “passive survivor” as described above; based on current knowledge, we boldly speculate that it may also act as an “active seeker” capable of dynamically correcting reprogramming errors. In contrast with the extra-embryonic lineages, EPI should undergo a profound genome-wide remodeling associated with establishment of pluripotency, a process considered dependent on its robust pluripotency regulatory network (e.g., *Oct4*, *Sox2*, and *Nanog*) and characterized by hallmark events such as global DNA demethylation^46,47^ and X-chromosome reactivation (XCR) achieved through the potent repression of *Xist*^48,49^. A compelling question arises: could this EPI-specific, powerful epigenetic remodeling mechanism, while establishing the naïve pluripotency, also concurrently purge the residual epigenetic aberrations derived from the somatic nucleus? If so, this would constitute a “secondary reprogramming” event specifically in the NT-EPI lineage to successfully reconstruct intact developmental pluripotency. This represents a highly intriguing hypothesis that warrants future investigation.

While our functional assays robustly demonstrate the full developmental potential of the NT-EPI lineage, a small number of DEGs departing from its FD compartment are still detected, consistent with the previous reports of EPI transcriptional defects in cloned blastocysts^50^. The NT-EPI has been further shown to be incapable of executing a successful naïve-to-primed transition due to the persistently aberrant activation of the Wnt signaling pathway, leading to the failure to correctly form the egg cylinder peri-implantation^50^. Rather than attributing these seemingly contradictory findings solely to incomplete reprogramming within the NT-EPI itself, we propose a reasonable and nonnegligible explanation: these aberrations may be extrinsically induced by its compromised extra-embryonic lineage neighbours. Given the well-established understanding of the precise and complex signaling crosstalk between embryonic and extra-embryonic lineages that crucially modulates EPI pluripotency and development^51^, it is highly probable that the extra-embryonic lineages with severe reprogramming disorder in cloned embryos fail to secrete signaling molecules or provide essential supportive cues normally, thereby inflicting potential “secondary damage” upon the molecular state and developmental trajectory of the EPI. For example, our scRNA-seq data reveal that the expression of *Dkk1*, encoding a key secreted inhibitor of the Wnt signaling pathway^52^, is significantly downregulated in NT-PrE cells (Figure S10), strongly suggesting a reduction of Wnt antagonist received by NT-EPI cells from surrounding microenvironment. This plausibly explains the previously reported persistent aberrant activation of Wnt signaling in NT-EPI, which results in defective peri-implantation development due to an inability to execute a successful naïve-to-primed transition^50^. Therefore, we posit that the detected molecular and developmental defects in NT-EPI should, at least partially, be attributed to suffering from the dysfunctional extra-embryonic components. This rationale underscores the necessity of holistically replacing the extra-embryonic lineages to establish an optimal “undisturbed” microenvironment for fully unleashing the true developmental potential of the NT-EPI, which ultimately proved to be equivalent to that of its FD counterpart.

In the present study, we adopted the induced lineages from E4.5 blastocysts as a surrogate, avoiding the technical challenge of physically dissecting late-stage blastocyst lineages and the concomitant risk of lineage cross-contamination, which ensures the accuracy of our reconstruction experiments. It is generally accepted that the lineage fates of EPI and PrE in E4.5 late-stage blastocysts are stable and irreversible^21,23,25^. Consistent with this, we observed that the iPrEs tenaciously maintained their lineage identity within reconstructed embryos, and both iEPIs and iPrEs correctly contributed to their respective embryonic and extra-embryonic tissues during the *in vivo* development of the tri-lineage aggregates. Surprisingly, a fraction of iEPIs underwent transdifferentiation towards extra-embryonic lineages, when solely aggregated with FD-4N embryos. This unconventional, seemingly “totipotent-like” behavior is unlikely to be an intrinsic property of E4.5 EPI, but is instead plausibly induced by the unique intercellular microenvironment formed within the iEPI/FD-4N reconstructed embryos. In this system, developmentally advanced iEPI cells are forced to aggregate with blastomeres from a much earlier developmental stage – 4-cell stage FD-4N embryos, which are speculated capable of creating a relatively undifferentiated signaling microenvironment that potentially induces fate reprogramming in the adjacent iEPI cells. In contrast, this special situation is not present in a naturally developing late-stage blastocyst, where the EPI lineage identity is stable post-specification. Nonetheless, our data unexpectedly reveal that the E4.5 EPI lineage still retains a latent degree of cellular plasticity, which can be unveiled under certain extreme non-physiological embryonic conditions.

Beyond the primary focus on validating the inherent full-term developmental potential of NT-EPI via tri-lineage aggregation, the present study, based on various lineage/embryo aggregation strategies, also demonstrates or reflects many aspects of lineage developmental characteristics and effects. Some comparisons of cloning birth rates among different aggregation groups highlight the significant impact of the PrE lineage on their full-term development (Table 1). For example, the NT-ICM/FD-4N group, despite possessing a presumably defective NT-PrE lineage, achieved a much higher birth rate than the NT-iEPI/FD-4N group (16.0% vs. 5.9%); conversely, when the latter was supplemented with a normal PrE lineage (NT-iEPI/FD-iPrE/FD-4N), the birth rate increased substantially (up to 20.5%). Both observations indicate that the presence of a PrE lineage at the onset of aggregation has a significant promoting effect on the full-term development of reconstructed cloned embryos. However, an excess of PrE cells may inversely have a detrimental effect on cloning efficiency, as indicated by the fact that the NT-ICM/FD-iPrE/FD-4N group, despite additionally including even a normal PrE lineage, exhibited a lower birth rate than the NT-ICM/FD-4N group (13.5% vs. 16.0%). Furthermore, corresponding to the aberrant transcriptome profile of NT-PrE that implies its functional defects, the birth rates obtained from NT-ICM/FD-4N and NT-ICM/FD-iPrE/FD-4N (16.0% and 13.5%, respectively) were both lower than that of the NT-iEPI/FD-iPrE/FD-4N group (20.5%), which included only an FD-PrE and no NT-PrE lineage at the start of aggregation. This may indirectly reflect the potential detrimental effect of NT-PrE on the full-term development of reconstructed cloned embryos. Far beyond the experimental data already obtained in this study, the innovative “Lego-like” induced lineage/embryo aggregation strategy we employed, when combined with lineage-specific transgenic fluorescence labeling and gene editing (knockout or overexpression), is envisioned to have broad applications in customizable studies on the tracing, interaction, and molecular regulatory mechanisms of lineages in embryonic development.

Despite the compelling functional evidence presented herein, our study still has several limitations. For instance, the omics assessment of the lineage reprogramming state is primarily based on the scRNA-seq data; while this can reliably indicate the reprogramming levels of lineages, the corresponding single-cell epigenomic profiles (e.g., whole-genome methylation or histone modifications) from the isolated NT-ICMs were unable to be obtained due to significant technical challenges. Second, although our “Lego-like” aggregation strategy allows for rigorous variable control, these artificially reconstructed embryos are speculated to possess an intercellular microenvironment different from that of natural development, which may exhibit a certain degree of negative impact on their development. This represents an intrinsic effect of the lineage/embryo aggregation strategies for both the experimental and control groups – an inevitable trade-off for achieving scientific validation. Finally, the deep molecular mechanisms driving the EPI-specific effective reprogramming remain to be elucidated in future studies.

In conclusion, our study uncovers a “lineage-specific asymmetric reprogramming” mode in SCNT blastocysts, where only the EPI lineage achieves effective reprogramming, constituting the deterministic developmental foundation for the generation of cloned animals. This finding not only provides a new theoretical basis for understanding SCNT reprogramming mechanisms, but also further defines the direction for improving cloning efficiency in the future – rescuing or replacing the defective extra-embryonic lineages, ideally in a holistic manner. Furthermore, the advanced lineage/embryo aggregation strategy we developed is, in itself, a powerful new tool with broad prospects for studying embryonic lineage development and interaction in both normal and aberrant contexts. From a broader perspective, SCNT can be viewed as an extreme stress model for investigating embryonic robustness, which here allows us to marvel at the extraordinary resilience embryos exhibit in the face of such severe epigenetic aberrations – a resilience ultimately concentrated within the embryonic (EPI) lineage destined to form life itself, thereby conferring and safeguarding the very possibility of cloned animal birth.

## Methods

### Mouse strains

The mouse strains used in this study include the wild-type C57BL/6J, DBA/2, ICR (CD-1) and FVB/N, and the C57BL/6J transgenic strains, *Nanog-GFP*, *Rosa26-CAG-tdTomato* (*R26-tdTomato*), and *CAG-GFP*. For the bulk RNA-seq and scRNA-seq of ICMs, FD blastocysts were obtained by mating C57BL/6J females with DBA/2 males, and NT blastocysts were generated using the cumulus cells and oocytes from B6D2F1 (produced by mating C57BL/6J females with DBA/2 males) females. For acquiring the (i)ICMs used for chimeric assessment in reconstructed blastocysts, FD 2-cell embryos were obtained by mating *Nanog-GFP;R26-tdTomato* homozygous females (produced by crossing *Nanog-GFP* and *R26-tdTomato* strains) with DBA/2 males, and NT 2-cell embryos were generated using the cumulus cells and oocytes from B6D2F1 *Nanog-GFP;R26-tdTomato* females (produced by mating *Nanog-GFP;R26-tdTomato* homozygous females with DBA/2 males). For acquiring the ICMs and iEPIs used for *in vivo* developmental potential assessment under lineage/embryo aggregation strategies, FD 2-cell embryos were obtained by mating *CAG-GFP* females with DBA/2 males, and NT 2-cell embryos were generated using the cumulus cells and oocytes from B6D2F1 *CAG-GFP* (produced by mating *CAG-GFP* females with DBA/2 males) females. For acquiring the FD-iPrEs used for *in vivo* developmental potential assessment under lineage/embryo aggregation strategies, FD 2-cell embryos were obtained by mating *R26-tdTomato* females with FVB/N males. For preparing tetraploid embryos, FD 2-cell embryos were obtained by intercrossing B6D2F1 (generating B6D2F2) or ICR. For embryo transfer, pseudo-pregnant ICR females were used as recipients.

### Fertilized embryo collection

Females of C57BL/6J (3–6 weeks old), B6D2F1 (8–12 weeks old) or ICR (3–8 weeks old) background received an intraperitoneal injection of 5–7.5 IU PMSG, followed 48 hours later by an equivalent dose of hCG. The females were then mated with males of corresponding strains as required by the experiments. The presence of a vaginal plug the next morning was designated as E0.5. At E1.5, 2-cell embryos were collected in HEPES-buffered CZB (HCZB) medium, washed, and transferred to preheated KSOM medium supplemented with amino acids (KSOM-AA) overlaid with mineral oil. Embryos were cultured at 37°C in an atmosphere of 5% CO₂ in air.

### Somatic cell nuclear transfer

B6D2F1 females (8–12 weeks old) used for SCNT experiments were superovulated by PMSG/hCG injection as described above, and MII oocytes were collected 14 hours post-hCG injection. The ampulla of the oviduct was punctured with a 30-gauge needle attached to a 1-mL syringe to release the cumulus-oocyte complexes (COCs). The COCs were treated with 0.1% hyaluronidase in HCZB medium for 3–5 min, and the denuded oocytes were then collected using a mouth-controlled pipette. Concurrently, the cumulus cells were collected in HCZB with 2.5% PVP and temporarily stored at 4℃, serving as nuclear donors. The residual hyaluronidase was removed from the oocytes by washing with HCZB, and the oocytes were then washed and cultured in CZB medium, overlaid with mineral oil at 37°C in 5% CO₂ in air.

The oocytes were enucleated using an 8–10 μm glass needle and maintained in CZB at 37°C. Piezoelectric pulses were applied to penetrate the oocyte membrane, facilitating the injection of a donor cumulus cell into an enucleated oocyte. Subsequently, the reconstructed oocytes were returned to CZB medium for 1-hour recovery, followed by activation for 5.5–6 hours in Ca²⁺-free CZB medium containing 10 mM SrCl₂, 5 ng/mL trichostatin A (TSA), and 5 µg/mL cytochalasin B (CB). All reconstructed embryos were then cultured in KSOM medium supplemented with 5 ng/mL TSA for another 3–4 hours, and finally maintained in KSOM medium with amino acids at 37°C under 5% CO₂ in air.

### Preparation of tetraploid embryos

FD 2-cell embryos of B6D2F2 or ICR background were subjected to electrofusion to obtain tetraploid embryos. Upon completion of the program, the embryos were removed from the apparatus and placed in fresh KSOM culture droplets, which were then transferred to an incubator for 30–60 min recovery. Thereinto, the resulting fused 1-cell embryos were considered tetraploid embryos. After thorough washing with KSOM medium, the 1-cell tetraploid embryos were transferred to fresh KSOM droplets for further culture.

### Preparation of induced lineages

For either the FD 2-cell embryos harvested from mated females or the NT 2-cell embryos developed *in vitro* after SCNT, 2i treatment was performed with KSOM medium supplemented with 1 µM PD0325901 and 3 µM CHIR99021 (termed 2i-KSOM) until the E4.5 late blastocyst stage to generate iEPI lineage (FD-iEPI and NT-iEPI, respectively), and F4H treatment was performed with KSOM medium supplemented with 1 µg/mL FGF4 and 1 µg/mL heparin (termed F4H-KSOM) until the E4.5 late blastocyst stage to generate iPrE lineage (FD-iPrE and NT-iPrE, respectively). These blastocysts post-induction were then treated with acidic Tyrode’s solution to remove the zona pellucida, followed by 10 washes in KSOM. The blastocysts were then transferred to rabbit anti-mouse serum (diluted 1:1 with KSOM) and incubated for 30 min at 37°C in 5% CO₂. After 10 washes in KSOM, the blastocysts were exposed to guinea pig serum (diluted 1:1 with KSOM) and incubated for 15 min at 37°C in 5% CO₂ until TE cell lysis was observed. The blastocysts were then recovered in KSOM for 30 min. Finally, lysed TE cells were mechanically removed by gentle pipetting with a 40–60 μm glass needle, and the isolated iICMs were then transferred to KSOM for temporary recovery and culture.

### Lineage/embryo aggregation and reconstruction

Aggregation plates were prepared using an aggregation needle to create depressions in the bottom of a 35-mm easy grip petri dish, as described previously^19,22^. For NT-iEPI/FD-iPrE/FD-4N aggregation strategy, two 4-cell FD-4N embryos after zona pellucida removement by acidic Tyrode’s solution were placed into the aggregation well, followed by transferring one NT-iEPI and one FD-iPrE into the depression. Upon the completion of aggregation operation, the aggregate was cultured in KSOM medium for 24 hours to obtain reconstructed blastocyst. The reconstructed blastocysts with significant chimerism of iICM cells (NT-iEPI and FD-iPrE) were selected by fluorescence microscopy and subsequently transferred into pseudo-pregnant recipients. For other tri-lineage aggregation strategies, the experimental procedures remained the same except for using different (i)ICM combinations. For dual-lineage (i)ICM/FD-4N aggregation strategies, one (i)ICM and two 4-cell FD-4N embryos were combined, and all other procedures were performed as described above.

### Embryo transfer

ICR females (8–16 weeks old) in good estrus were paired with vasectomized males. The day following plug detection was designated as E0.5. At E2.5 of these pseudo-pregnant recipients, the NT embryos or reconstructed embryos cultured *in vitro* to the blastocyst stage were transferred into the uterus of pseudo-pregnant recipients. Generally, fetuses were delivered by cesarean section at E18.5, and data related to fetuses and placentae were recorded and analyzed. Additionally, nursing ICR mice were prepared in advance for fostering the newborn pups.

### Single-cell dissociation of inner cell mass

FD or NT embryos were cultured *in vitro* to the E4.5 late blastocyst stage. The zona pellucida was removed by brief exposure to acidic Tyrode’s solution. Immunosurgery was then performed to lyse the outermost TE cells using anti-mouse serum and guinea pig complement. The lysed TE cells surrounding the ICM were mechanically aspirated and discarded using glass needles of various diameters (100–120 μm and 40–50 μm). Subsequently, the ICM was digested with trypsin-EDTA in calcium-free medium for 5–10 min at 37°C. Single-cell dissociation was then performed using a 40–60 µm glass needle and a micromanipulator (Movie S1). The dissociated single cells were washed 3 times in 0.1% BSA solution, and individually collected for subsequent scRNA-seq.

### RNA library preparation and sequencing

The single cells isolated from the ICMs of E4.5 FD or NT blastocysts were individually transferred into tubes containing 200 µL lysis buffer by a mouth-controlled pipette. After lysis at 72°C for 3 min, reverse transcription mixture was added into each tube, followed by incubation at 25°C for 5 min, 42°C for 1 hour, and 50°C for 30 min, with a final inactivation step at 70°C for 10 min. The resulting cDNA samples were subsequently amplified, and purified using AMPure XP beads. The DNA abundance was enhanced through PCR using the Nextera XT DNA sample preparation kit. According to the manual of the KAPA HyperPrep Kit, single-cell RNA-seq libraries were created, and then sequenced on the Illumina Hiseq platform. For the preparation and sequencing of bulk RNA-seq libraries, the isolated E4.5 (i)ICMs were individually transferred into tubes containing 200 µL lysis buffer, all other procedures were the same as described above.

### RNA-seq data processing and analysis

For both the scRNA-seq and bulk RNA-seq data, raw reads were trimmed using Trim Galore (v0.6.10) with default parameters to remove low-quality bases and adapter sequences, and then aligned to the mouse reference genome (mm10) using Salmon (v0.14.1). Gene expression levels were quantified by the normalization of transcripts per million mapped fragments (TPM). DEGs between groups were identified using the Wilcoxon rank-sum test with FDR correction (*p* < 0.05). Prior to DEG analysis, scRNA-seq data were processed to assign EPI or PrE lineage identity to each cell based on the expression of established lineage-specific marker genes^33,50,53^ (Figure S1A, B), and the classification was further validated by examining additional reported lineage markers^33^ (Figure S1C, D).

### Immunofluorescence staining

Reconstructed blastocysts were fixed in 4% PFA at room temperature for 30 min, followed by washing 3 times with PBS containing 0.5% BSA. Subsequently, the embryos were permeabilized using 0.1% Triton X-100 in PBS (PBTX) for 30 min, and washed 3 times by PBS. Transferred to 3% BSA in PBS, the embryos were blocked at room temperature for 1 hour, and then incubated with the primary antibody diluted in the 3% BSA (1:200) overnight at 4°C. The embryos were washed 3 times by PBS, followed by incubation with the secondary antibody diluted in the 3% BSA (1:500) for 1–2 hours at room temperature. The nuclei were then stained with DAPI for 5 min, and the embryos were washed 3 times with PBS before being observed and photographed under the LSM880 confocal microscope. The primary antibodies used in the present study include rabbit anti-Nanog (Abcam) and goat anti-Gata6 (R&D Systems). The corresponding secondary antibodies used in the present study were purchased from Thermo Fisher Scientific.

### Frozen sectioning

Placental tissues collected from E18.5 mouse conceptuses were washed briefly in cold PBS to remove excess blood, and fixed in 4% paraformaldehyde (PFA) at 4°C overnight. Fixed tissues were cryoprotected in graded sucrose solutions (15% and 30% in PBS) at 4°C, embedded in optimal cutting temperature (OCT) compound, and frozen at −80°C. Frozen blocks were sectioned at 7 µm thickness using a cryostat (Leica CM1950), mounted onto glass slides, air-dried for 30 min at room temperature, and stored at −80°C until use.

### H&E staining

Frozen sections were removed from −80°C storage, equilibrated to room temperature for 10–15 min, and briefly rinsed in PBS to remove residual OCT compound. Sections were refixed in 4% PFA for 10 min, followed by 4 washes with deionized water. Then, sections were stained with hematoxylin for 3 min, rinsed in tap water, differentiated in 1% acid alcohol, and counterstained with eosin for 30 sec. After dehydration through graded ethanol and clearing in xylene, sections were mounted with neutral balsam and observed under a light microscope. Images were captured using a digital camera attached to the microscope.

### Statistical analysis and figure preparation

Statistical analyses and visualization were performed using GraphPad Prism 9.0. Visualization of DEG analyses was generated using the ggplot2 package in R. Schematic diagrams were created and image assembly was completed using Adobe Photoshop and Illustrator.

## Supporting information

Demonstration of dissociating ICM micro-masses into single cell

## Declarations

### Ethics statement

Mouse care and all the experimental procedures were conducted in compliance with the guidelines of the Institutional Animal Care and Use Committee (IACUC) of the Kunming Institute of Zoology, Chinese Academy of Sciences. The approval number for all the contents of this research is IACUC-RE-2024-01-006.

### Data availability

All data supporting the findings of this study are available within the paper and its Supplementary Information. Additional analysis data are available from the corresponding author upon reasonable request.

### Competing interests

The authors declare that they have no competing interests.

### Authors’ contributions

J.L. conceived this project. G.L, J.L., M.H., W.B., H.X., S.W., J.X., Y.T., Y.Z., Y.C., L.R., J.L., and J.D. performed the experiments. L.L. performed sequencing data analysis. Z.M. provided guidance. J.L., W.B., G.L., M.J., and C.Z. drafted the manuscript. All authors read and approved the final manuscript.

## Acknowledgements

We sincerely thank Prof. Jinsong Li, Maria Elena Torres-Padilla, Ping Zheng, and Shuhui Bian for critical reading of the manuscript. This work was supported by the National Natural Science Foundation of China (31970823 and 32270862).

## Supplementary Information

**Figure S1.**
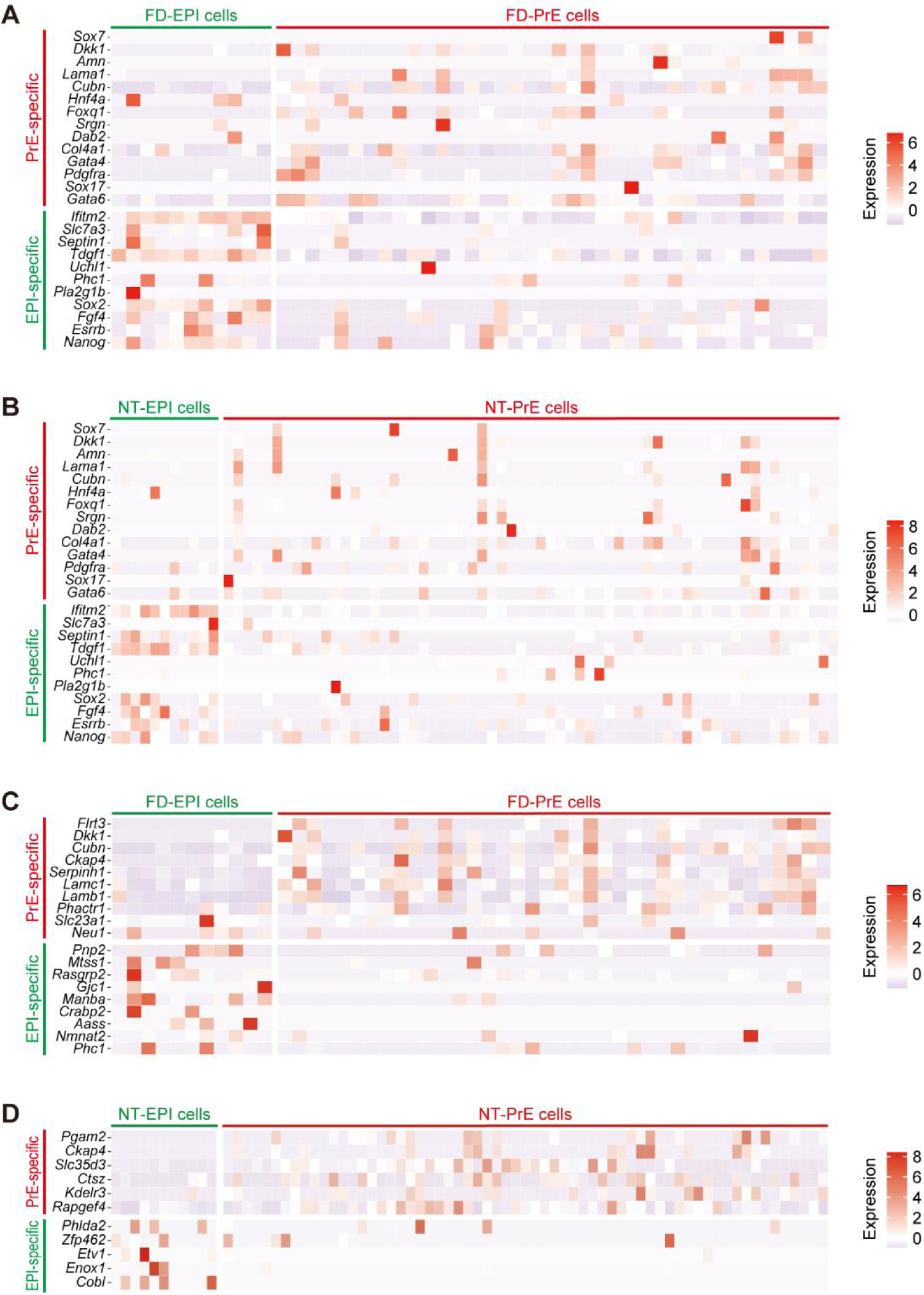
Identification of EPI and PrE lineage cells in the scRNA-seq of E4.5 FD-ICMs and NT-ICMs. (A, B) Expression heatmaps of lineage-specific marker genes used to identify the EPI and PrE lineage cells in the scRNA-seq of E4.5 FD-ICMs (A) and NT-ICMs (B). (C, D) Expression heatmaps of additional lineage-specific marker genes used to further confirm the EPI and PrE lineage cells identified in the scRNA-seq of E4.5 FD-ICMs (C) and NT-ICMs (D).

**Figure S2.**
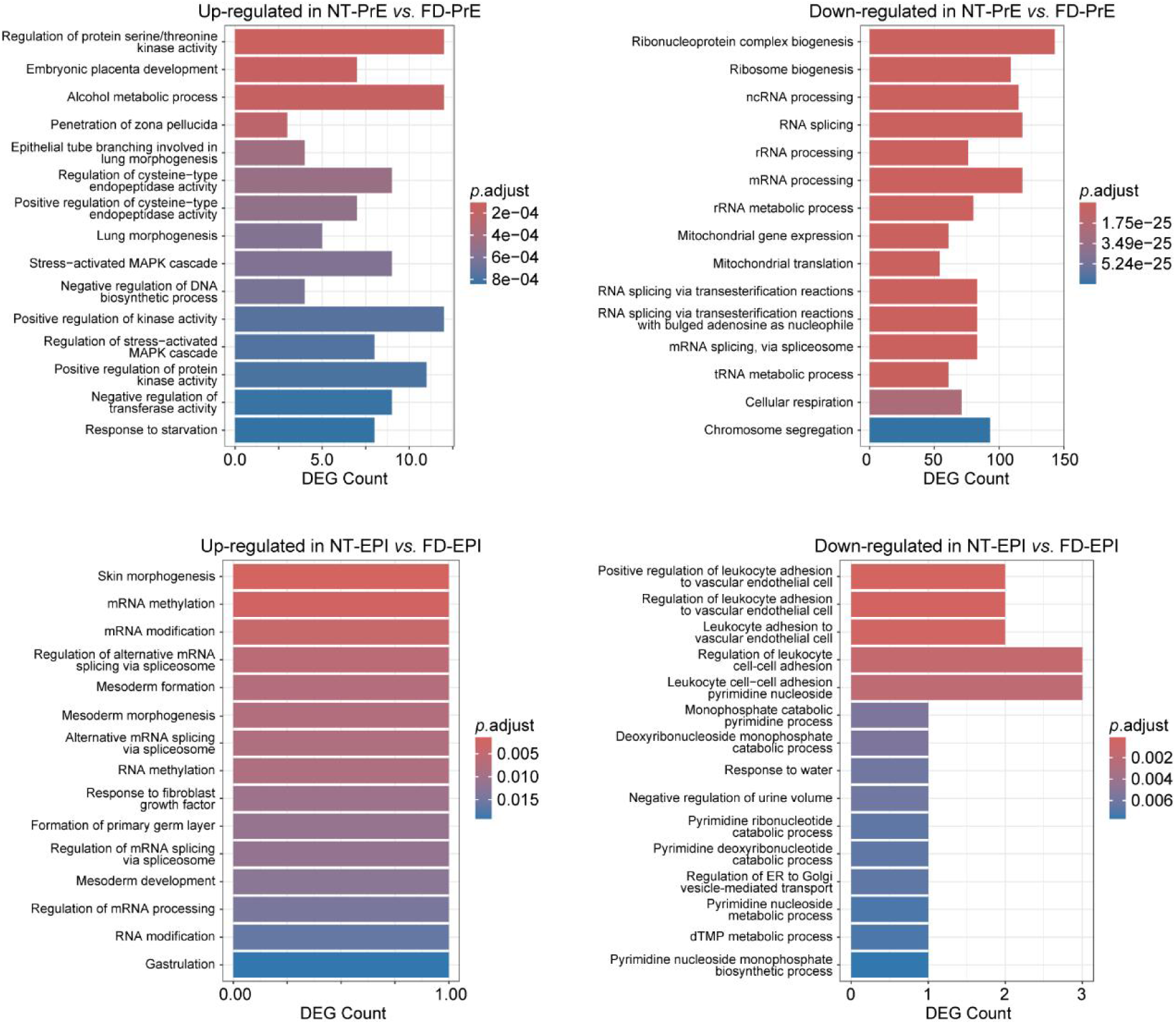
GO enrichment analyses on the DEGs in E4.5 NT-PrE *vs*. FD-PrE and NT-EPI *vs*. FD-EPI. The up-regulated and down-regulated DEGs were analyzed separately. Representative BP terms are shown for each analysis.

**Figure S3.**
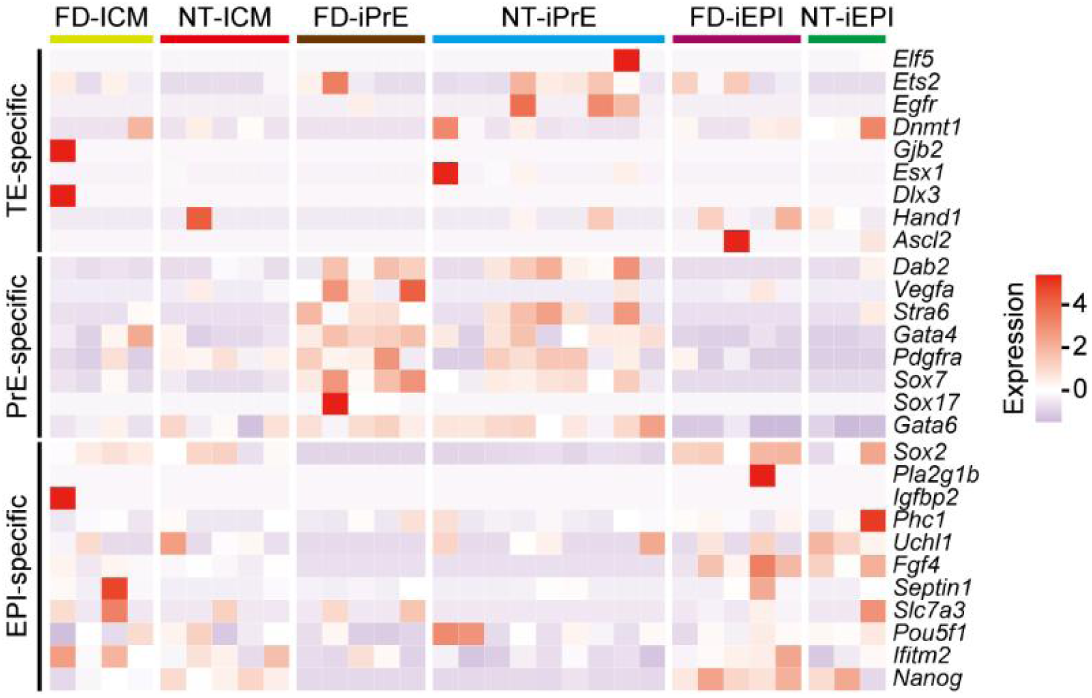
Expression pattern of lineage-specific marker genes in the bulk RNA-seq of each (i)ICM type.

**Figure S4.**
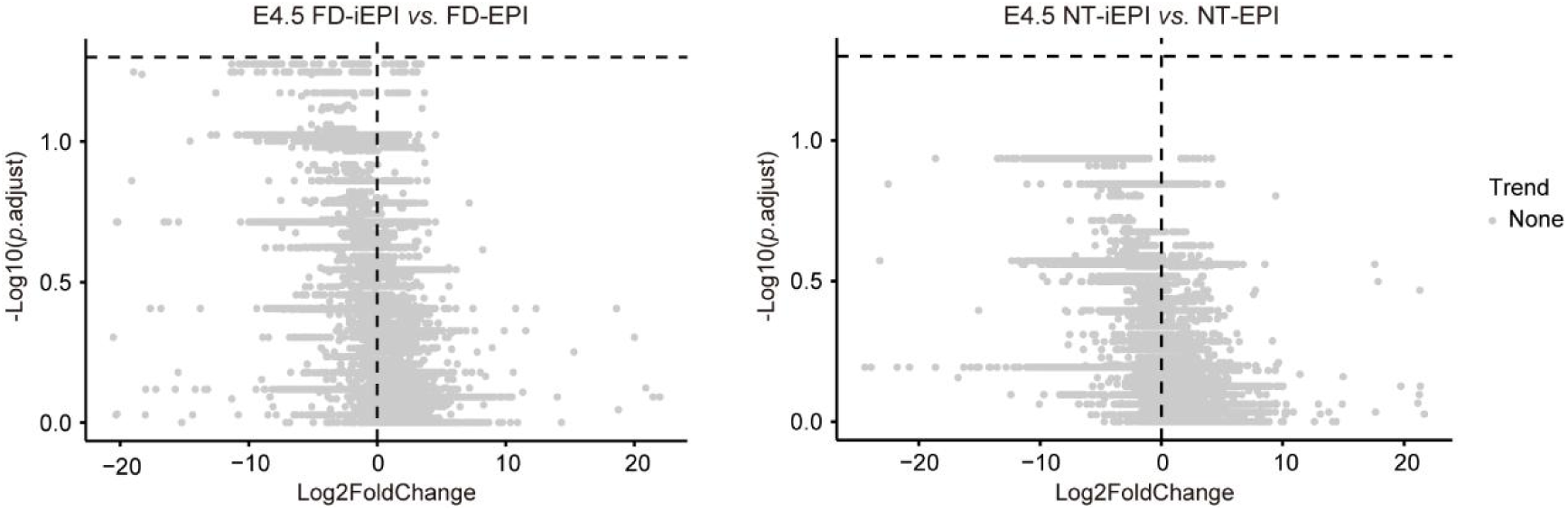
DEG analyses in E4.5 NT-iEPI *vs*. NT-EPI and FD-iEPI *vs*. FD-EPI. The bulk RNA-seq data of E4.5 NT-/FD-iEPIs and the transcriptional profiles of EPI lineage cells in the scRNA-seq of E4.5 NT-/FD-ICMs were correspondingly utilized for the DEG analyses (*p*.adjust < 0.05).

**Figure S5.**
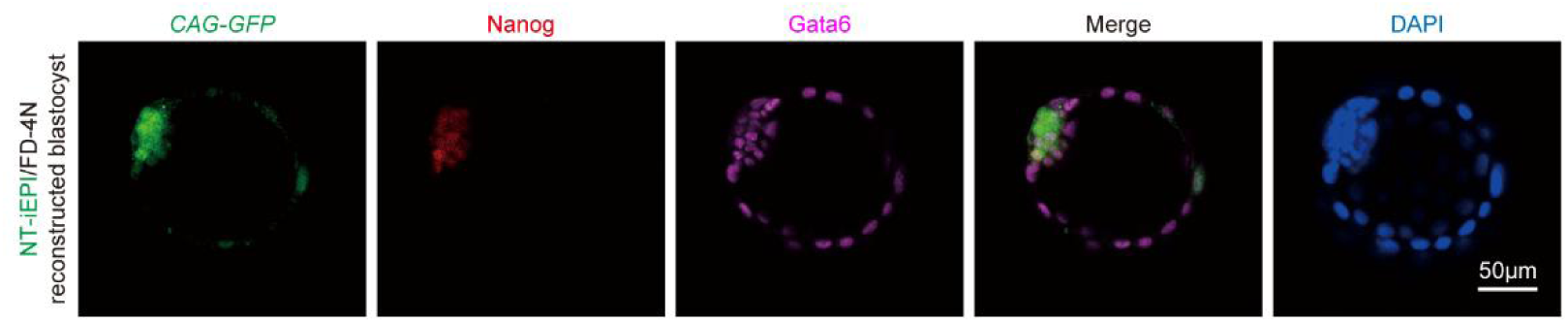
Immunofluorescence staining of NT-iEPI/FD-4N reconstructed blastocyst for *Nanog* and *Gata6* expression. The NT-iEPI(-derived) cells were of *CAG-GFP* transgenic background. The reconstructed blastocysts for immunofluorescence staining (*n* = 13) were collected after 48-hour *in vitro* culture post-aggregation.

**Figure S6.**
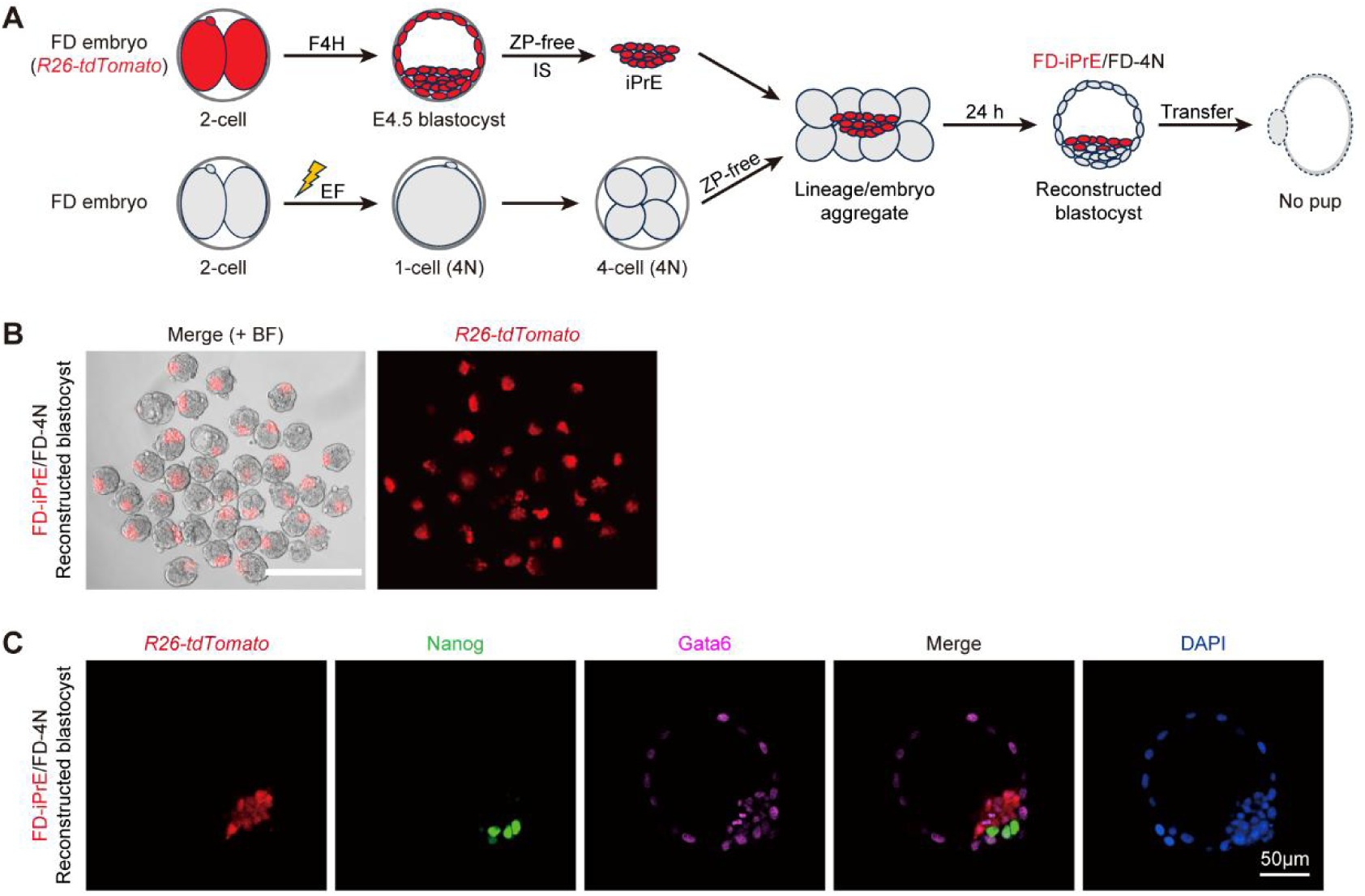
Assessment of the *in vivo* developmental potential of FD-iPrE under tetraploid complementation. (A) Schematic illustration of strategy for assessing the *in vivo* developmental potential of FD-iPrE under tetraploid complementation. The *R26-tdTomato* FD embryo was treated with F4H from the 2-cell stage until the E4.5 blastocyst stage; then, the resulting FD-iPrE was isolated via removal of zona pellucida and immunosurgery, and aggregated with two 4-cell FD-4N embryos; after 24-hour *in vitro* culture, the reconstructed blastocyst developed from the aggregate was subsequently transferred to assess its *in vivo* developmental potential. ZP, zona pellucida; IS, immunosurgery; EF, electro-fusion. (B) Fluorescence microscopy of the FD-iPrE/FD-4N reconstructed blastocysts after 24-hour *in vitro* culture post-aggregation. Scale bar = 50 μm. (C) Immunofluorescence staining of FD-iPrE/FD-4N reconstructed blastocyst for *Nanog* and *Gata6* expression. The FD-iPrE(-derived) cells were of *R26-tdTomato* transgenic background. The reconstructed blastocysts for immunofluorescence staining (*n* = 17) were collected after 48-hour *in vitro* culture post-aggregation.

**Figure S7.**
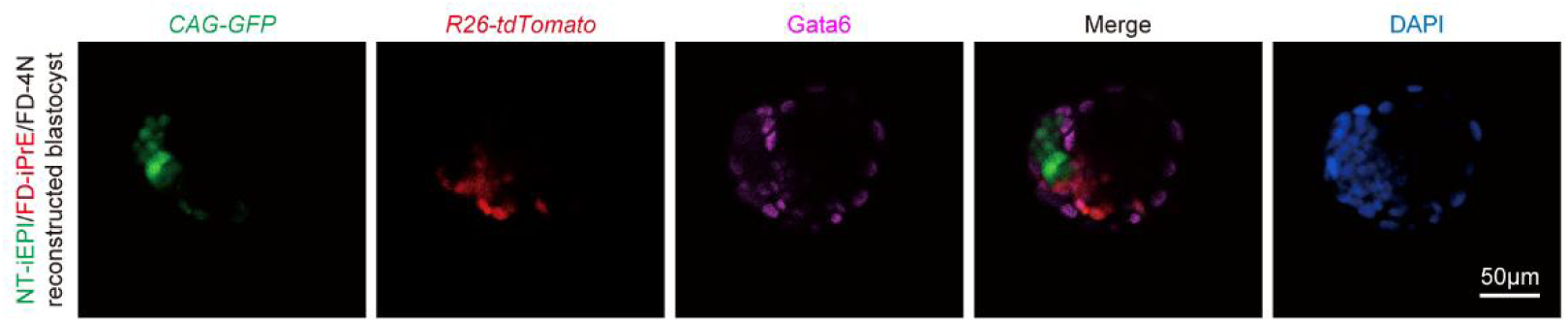
Immunofluorescence staining of NT-iEPI/FD-iPrE/FD-4N reconstructed blastocyst for *Gata6* expression. The NT-iEPI(-derived) and FD-iPrE(-derived) cells were of *CAG-GFP* and *R26-tdTomato* transgenic background, respectively. The reconstructed blastocysts for immunofluorescence staining (*n* = 9) were collected after 48-hour *in vitro* culture post-aggregation.

**Figure S8.**
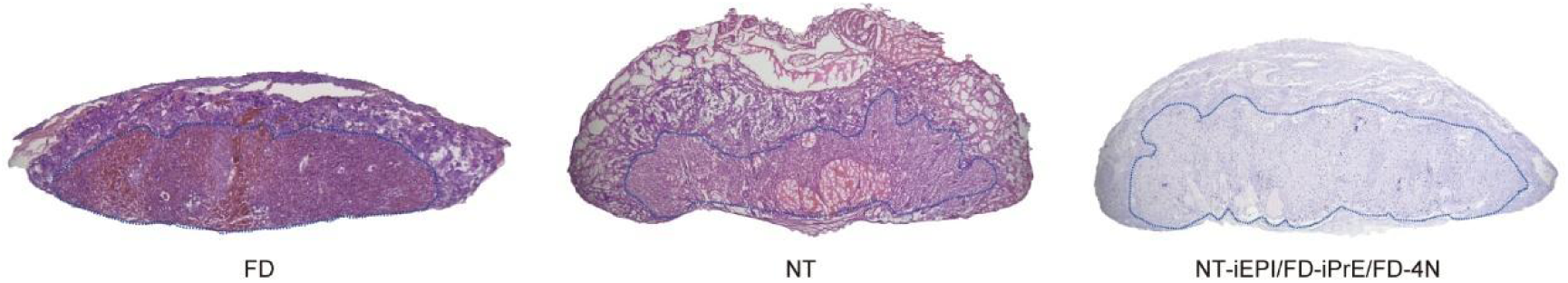
Histological sections with H&E staining show the phenotypic rescue of E18.5 placenta generated by NT-iEPI/FD-iPrE/FD-4N strategy. The sections of FD and NT placentae serve as the normal and pathological controls, respectively. For each sample, the labyrinth layer is outlined by the blue line, with the spongiotrophoblast layer located above.

**Figure S9.**
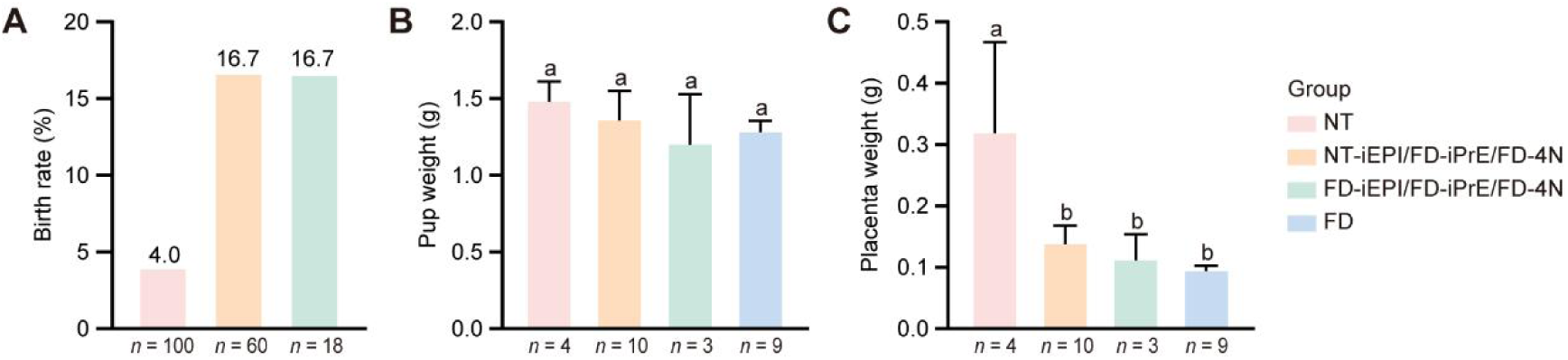
Statistical analysis of the neonatal data obtained from repeated tri-lineage embryonic aggregation strategies. Statistical aspects include the birth rate (A), body weight (B), and placenta weight (C) of the neonatal pups. The FD and NT groups serve as the normal and pathological controls, respectively. Statistical significance was determined by Student’s t-test. Groups with different letters (a, b) indicate significant differences (*p* < 0.05), while groups sharing the same letter are not significantly different (*p* > 0.05).

**Figure S10.**
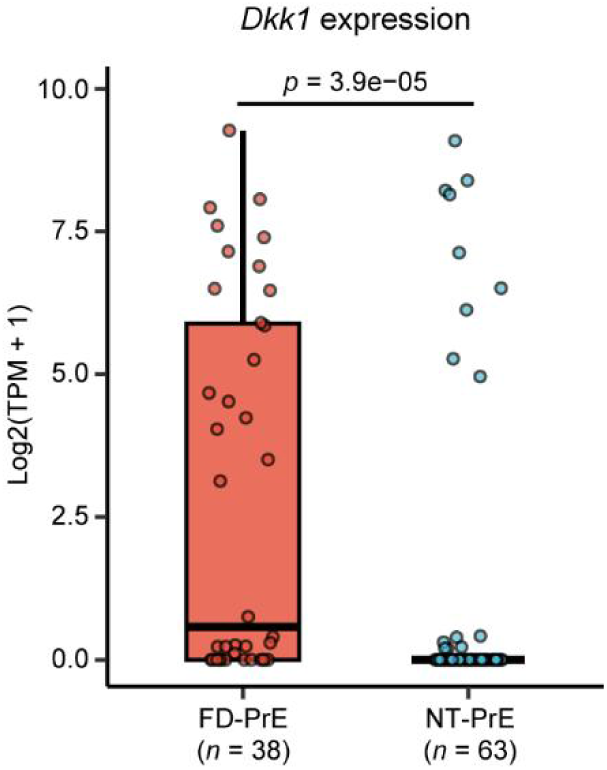
*Dkk1* expression levels in the FD-PrE and NT-PrE lineages. The gene expression data was acquired from the transcriptional profiles of PrE lineage cells in the scRNA-seq of E4.5 FD-ICMs and NT-ICMs. Statistical significance was determined by Wilcoxon rank-sum test.

**Movie S1. Demonstration of dissociating ICM micro-masses into single cells.**

## Notes

### Competing Interest Statement

The authors have declared no competing interest.

